# Astrocytic regulation of synchronous bursting in cortical cultures: from local to global

**DOI:** 10.1101/870915

**Authors:** Ravi Kumar, Yu-Ting Huang, Chun-Chung Chen, Shun-Fen Tzeng, C. K. Chan

## Abstract

Synchronous bursting (SB) is ubiquitous in neuronal networks. It is known for a long time that SB is driven by glutamatergic neurotransmissions but its underlying mechanism is still unclear. Recent studies show that local glutamate recycle by astrocytes can affect neuronal activities nearby. Since SB is independent of network structure, it is conceivable that the local dynamics might also be the origin of SB in networks. We investigated the effects of local glutamate dynamics on SBs in both cultures developed on multi-electrode array (MEA) systems and a tripartite synapse simulation model. In our experiments, local glutamate recycle dynamics are studied by pharmacologically targeting the astrocytic glutamate transporters (GLT-1), while neuronal firing activities and synaptic glutamate level are simultaneously monitored with MEA and glutamate sensor (iGluSnFR) expressed on surface of astrocytes respectively. We found SBs to be synchronized with glutamate transients and the manipulation of local glutamate dynamics can indeed alter the global properties of the SBs. Detailed simulation of a network with astrocytic glutamate uptake and recycle mechanisms conforming with the experimental observations revealed that astrocytes function as a slow negative feedback for the neuronal activities in the network. With this model, SB can be understood as the alternation between the positive and negative feedback in the neurons and astrocytes in the network respectively. An understanding of this glutamate trafficking dynamics is of general application to explain disordered phenomena in neuronal systems, and therefore can provide new insights into the origin of fatal seizure-like behavior.

**Significance:** Synchronous bursting (SB) is a hallmark of neuronal circuits. Contrary to the common belief that the SB is governed mainly by neuron-neuron interactions, this study shows that SBs are orchestrated through a generic neuron-astrocyte tripartite interactions. These interactions, identified as glutamate uptake and recycle processes in astrocytes, control the excitability of neuronal networks and shape the overall SB patterns. Our simulation results suggest that astrocytes traffic more glutamate than neurons and actively regulating glutamate proceedings around synapses. A bipartite synapse is a good approximation of a tripartite synapse provided that astrocyte-dependent glutamate content is taken into account. Our findings provide key insights into the ubiquity of SB and the origin of fatal seizure-like behavior in brain arising from astrocytic malfunction.

## Introduction

Synchronous firings of neurons are frequent in brains which can emerge as patterns of low frequency spiking or synchronous bursting (SB), where the SBs are believed to produce functions of a network (Arenas et al., 2008). Although astrocytes and neurons both constitute our brains, the SBs are commonly considered to result from neuron-neuron interactions alone (von Krosigk et al., 1993; Sirota et al., 2008; Penn et al., 2016). However, astrocytes are known to modulate nearby neuronal activities by forming tripartite synapses and exhibiting calcium-excitability in response to surrounding neuronal activities (Araque et al., 1999; Pascual et al., 2005). Therefore, a possible astrocytic mechanism for SB cannot be ruled out. Recent findings show that astrocytes play an important role in shaping neuronal firings by regulating synaptic glutamate via GLT-1 glutamate transporters (Murphy-Royal et al., 2015); providing evidence to a possible astrocytic mechanism in SB. However, it still remains unclear how the local properties of a synapse relate to the SB phenomenon, which is a network behaviour.

Since SBs are ubiquitous in both structured brains (Axmacher et al., 2006; Hutchison et al., 2004) as well as randomly connected cortical cultures (Wagenaar et al., 2006; Adams et al., 2011), local astrocytic modulation of neuronal firings possibly could be the basic mechanism for SB generation in a network, independent of connection topologies. In the SB phenomena, the neuronal system can be described to alternate between two states; ‘active state’ (during an SB) and ‘dormant state’ (time between SBs). Intuitively, the pools of ready-to-be-released glutamate in the neurons must deplete during the “active” state and undergo recovery in the “dormant” state. Our view of SB mechanism is that this alternation between pool depletion and replenishment is intimately related to the recycling of glutamate in the tripartite synapses formed by neurons and astrocytes in the network.

In a traditional bipartite synapse model, as proposed by Tsodyks, Uziel and Markram (TUM), neuronal excitability is characterized by recycling of a conserved amount of synaptic resources (Tsodyks et al., 2000). In this TUM model, synaptic resources (i.e. glutamate) are recycled through three states: recovered, active, and inactive. It is the recovered state which provides excitability to neurons. However, experiments have shown that released glutamate are uptaken by both neurons and astrocytes, and 80-90% of the released glutamate are uptaken by astrocytes (Danbolt et al., 1998) which were not considered in the TUM model. Recently, astrocytic GLT-1 was found to regulate synaptic currents (Murphy-Royal et al., 2015) and the period of SBs Huang et al. (2017b). It is also known that glutamate transported into the astrocytes get converted into glutamine and transported to neurons as precursors for glutamate, viz. glutamate-glutamine cycle (Danbolt et al., 1998; Hertz et al., 1999). Since the timescale of astrocytic glutamate recycling pathway is much longer than the presynaptic neuronal pathway, the recycle timescale difference should generate multiple timescales of glutamate depression. To account for the multiple timescales in SB, (Volman et al., 2007) extended the TUM model by introducing a hypothetical ‘super-inactive’ state for glutamate. In our view, this super-inactive state could just be the glutamine in the astrocyte, consistent with the findings of (Tani et al., 2014; Bacci et al., 2002) which show direct evidence on the necessity of synaptically localized glutamate-glutamine cycle for glutamate synthesis at excitatory terminals. These lines of reasoning strongly support that SB could be the consequence of astrocyte-mediated glutamate regulation in response to neuronal dynamics.

We hypothesized that SBs in neuronal networks arise from the local effects of the glutamate recycling in the tripartite synapses. We tested this hypothesis by both experiments and numerical simulation. In experiments, glutamate transporters were targeted in cortical cultures developed on multi-electrode array (MEA). Combining astrocyte-specific fluorescent probes with MEA, responses from neurons and astrocytes were recorded simultaneously. To test the validity of astrocytic regulation of SB, we developed a tripartite synapse model to include astrocytes in the glutamate recycling process. Results from both our experiments and simulations were found consistent with our hypothesis. Our novel finding suggests that the forms of SB are governed by the amount of glutamate residing in astrocytes. This later fact can be used to understand the epileptic form of firings (SB) in GLT-1 dysfunction (Rothstein et al., 1996; Coulter and Eid, 2012).

## Materials and Methods

All samples from animals were prepared according to the guidelines approved by Academia Sinica IACUC (Protocol: 12-12-475). All pharmacological experiments and simultaneous recordings of MEA and iGluSnFr-glutamate imaging / GCaMP6f-calcium imaging were performed on cultures (age *>* 20 DIV) inside home-made incubation chamber maintained at 5% CO_2_ - 95% air at 37°C.

### Cell culture

Cell culture protocol was adapted from Potter and DeMarse (2001). Briefly, neocortical tissues were obtained from postnatal mice pups (P0) FVB/NJ-wild type under sterile conditions. Pieces of dissected cortices were first enzymatically dissociated at 37°C with 0.125% trypsin solution for 10 minutes and then mechanically triturated with sterile and fire-polished Pasteur glass pipette. From this cell-suspension, cells were plated on 4-well MEA probes at 2000 cells/ mm^2^. The surfaces of the plating areas were previously coated with 0.1% polyethylenimine(PEI) and laminin for at least two hours. Cells in the remaining suspension were seeded into flasks and cultured simultaneously for obtaining conditioned media (CM). All the cultured cells were maintained in NeuroBasal-A medium (Invitrogen) supplemented with GlutaMax and B27, inside a humidified incubator with 5% CO_2_ - 95% air at 37°C. To prevent evaporation and reduce chances of contamination culture wells were always sealed with home-made teflon membrane caps. From each well (total volume capacity - 250*µ*L), half (125*µ*L) of the maintenance media was replaced with fresh maintenance media in every three days.

### Adeno-associated virus infection

At the age of 7 days *in vitro*(DIV), selected developing cortical networks were infected with AAV.GFAP.iGluSnFR to express GFAP-promoted glutamate sensors in astrocytes. The virus pENN.AAV.GFAP.iGluSnFr.WPRE.SV40 was a gift from Loren Looger (Addgene plasmid cat.# 98930). At the same time, few other growing cortical networks were virally infected with gfaABC1D-driven (similar to GFAP promoter) cyto-GCaMP6f to express calcium sensors in astrocytes. The virus pZac2.1 gfaABC1D-cyto-GCaMP6f was a gift from Baljit Khakh (Addgene plasmid cat.# 52925). Viral infections were performed at upto 20×10^9^ viral particles as final concentration in 200*µ*L volume per well. 24 hours post-infection, samples were washed with conditioned media and maintained in the usual manner for more than two weeks before performing any fluorescence imaging or pharmacological experiment.

### Pharmacology

All the drugs used in this study were purchased from Tocris Bioscience, United Kingdom unless otherwise noted. To obtain more reproducible results from drug experiments, the following protocol was adapted from Keefer et al. (2001). Briefly, prior to any pharmacology experiment the maintenance media of test sample well was completely replaced and bathed with 200*µ*L CM (pooled from flask cultures). Following a 30 minutes of stabilization, a baseline activities of the test sample was recorded for the next 30 minutes. Test drug was then added (maximum volume upto 2*µ*L), gently mixed by pipetting through micropipette. After drug addition, cultures were again left to stabilize for the next 10 minutes. Influence of test drugs on network firing activities were usually visible within five minutes of addition. Steady state response of the network was then recorded. Drugs were washed with the same CM three times within a span of five minutes. After another 30 minutes of stabilization, recovered activities were recorded. The drugs and their final concentrations used:Dihydrokainic acid (DHK, GLT-1 inhibitor, 200*µ*M, cat. no.#0111);DL-threo-beta-Benzyloxyaspartate (DL-TBOA, non-selective EAAT inhibitor, 10*µ*M, cat. no.#1223);GT949 (positive allosteric modulator of EAAT2, 10*µ*M, cat. no.#6578);Bicuculline methiodide (Bic, GABA antagonist, 10*µ*M, Abcam, cat. no.#ab120108;Riluzole hydrochloride (Na^+^ channel inhibitor, 10*µ*M, cat. no.#0768)).

### Electrophysiological recording

Cultures were developed on four-well multi-electrode planar arrays purchased from Qwane Biosciences SA, Switzerland (MEA-60-4well-PtB). Each of the four wells were embedded with 14 platinum black electrodes laid out in a square grid. The electrodes (diameter 30 *µ*m each) were separated by 200 *µ*m. Electrical signals from active cortical networks were sampled at 20kHz, amplified using MEA-60 pre-amplifier and digitized using MC Card PCI board in vitro MEA-system (MEA1060-Inv-BC, Multi Channel Systems MCS GmbH, Germany). Electrode configuration settings and data acquisition were performed with MEA Select and MC Rack softwares (both MultiChannel Systems, Germany) respectively. For long duration recording (upto several hours), an incubation chamber was constructed around the MEA system consisting of a temperature controller maintaining 37°C and CO_2_ supply. Pharmacological experiments were performed on mature samples (*>*20 days *in vitro*) which showed stable array-wide synchronous burst events.

### Fluorescence Microscopy

iGluSnFr or GCaMP6f imaging was performed inside a custom home-made incubation chamber built on a phase contrast microscope platform for maintaining 5% CO_2_ - 95% air at 37°C. For obtaining higher sensitivity and good signal to noise ratio for longer continuous recording (up to 15 minutes per session), a CCD camera (Prosilica GE680, Allied Vision, Canada) was coupled with an image intensifier(II18, Lambert Instruments, the Netherlands). For epifluorescence illumination, a mercury arc lamp illuminator (X-cite, Lumen Dynamics, Canada) was used. Calcium/ glutamate fluorescence signals from test samples were captured with 640×480 resolution at 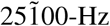 sampling rate on 20X objective, covering an area of ≈0.3mm^2^.

### Immunocytochemistry

After experiments, selected samples were fixed with 4% PFA for immunocytochemistry. Neuronal soma and dendrites were identified with microtubule-associated protein 2 (MAP2, Sigma, cat no.#M4403).Neuronal axon projections and their connectivity was identified with neuron-specific class III *β* -tubulin (Tuj1, GeneTex, cat no.# GTX85469) primary antibody. For identifying astrocytes and glutamate transporter (GLT-1) expression, glial fibrillary acidic protein (GFAP, Sigma, cat no.#G3893) and GLT-1 marker antibody (Invitrogen, cat no.#701988) were used respectively. Nuclei of all cell types were identified with DAPI staining. Confocal imaging of immuno-stained samples were performed at 10X/20X magnification using inverted confocal microscope (ZEISS LSM 880 system, Germany).

### Analysis

Data analysis was performed using MATLAB (MathWorks), ImageJ (National Institutes of Health - public domain) and Prism (GraphPad) software.

#### MEA data preprocessing

Data acquired from Multichannel System recordings, were imported into MATLAB for further processing and analysis. Before spike detection, raw data was pre-processed through a Butterworth 2^nd^ order high-pass filter, with cutoff frequency (200 Hz) to remove low frequency line and other system generated noise. Data presented here are not spike sorted. For each well, spike times from all the 14 channels were detected using Precise Timing Spike Detection (PTSD) algorithm (Maccione et al., 2009) and arranged in ascending order (*t*_1_, *t*_2_, *…, t*_*N*_).

#### Synchronous burst detection

Synchronous bursts were detected using the method described in Huang et al. (2017a). Briefly, firing rate time histogram of all channels combined was generated with a bin size of 10 ms. For every network its maximum firing rate (*R*_max_) was determined. Two threshold criteria (a lower *ER*_max_ followed by an upper Δ*R*_max_) were required to be satisfied, where *E*=0.04 and Δ=0.2 typically. The lower threshold decided whether a network’s active state is initiated. Starting of a burst is registered at *t*_*s*_ once the network becomes active at *t*_*s*_ and remains active until its summed spike rate reaches its upper threshold. Burst end *t*_*e*_ is registered when the network becomes inactive and remains inactive at least for a duration of 1 second (*τ*_rest_). After post-hoc manual inspection, *E*, Δ and *τ*_rest_ were slightly tuned whenever it was required to increase the efficiency of SB detection. To compare the level of synchronous bursting activities across different networks *SB index* was defined, as the ratio of sum of spikes detected within all SBs to the sum of all spikes detected array-wide, i.e.,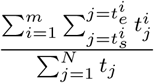 where *t*_*j*_ is a detected spike time and *m* is total number of SBs detected. An SB index of 0.5 indicates that 50% of array-wide spiking activities occurred in the form of SBs.

#### Fluorescence imaging data preprocessing and analysis

An instantaneous fluorescence intensity (*F*_*t*_) - time series was generated for each recording using *Plot Z-axis Profile*, ImageJ plugin. For each frame, *F*_*t*_ was calculated by averaging its pixels values. Further analyses were carried out in MATLAB. Drifts and photo-bleaching trend lines were removed using a polynomial curve fitting method, where a fitted curve was subtracted to remove any trend. Change in fluorescence Δ*F/F* was then defined as (*F*_*t*_ − *F*)*/F*, where *F* represented the mean instantaneous fluorescence intensity of the recording.

### Tripartite synapse TUMA Model

Our tripartite TUMA model extends on the short-term synaptic depression model described by Tsodyks, Uziel and Markram (Tsodyks and Markram, 1997; Tsodyks et al., 2000), hence referred to as TUM model, to consider the effects of astrocyte. The TUM model implements three states of synaptic resource trafficking with neuronal transmitters represented by three variables: *X, Y*, and *Z*, corresponding to the fractions of the resource in the recovered, active, and inactive states respectively. Assuming synaptic resource is a conserved quantity, the relation *X* + *Y* + *Z* = 1 will hold. Here we identify synaptic resources as glutamate. Upon spike arrival at time *t* = *t*_spk_, the transitions of glutamate between the three states are given by the equations,

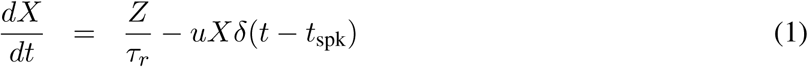

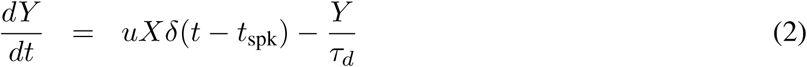

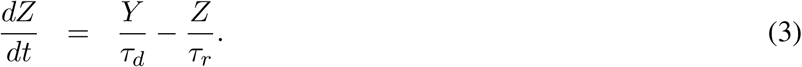

Equations 1–3 describe the transportation, transformation, and conservation of the synaptic glutamate under the spiking of the pre-synaptic neuron, with *u* being the facilitation factor provided from calcium influx. The relevant time scales are *τ*_*r*_, as the recovery time for the transformation from *Z* to *X*, and *τ*_*d*_, as the decay time from *Y* to *Z* are phenomenological. This model is shown to well reproduce the synaptic responses between pyramidal neurons (Tsodyks and Markram, 1997).

When an action potential arrives at the TUM synapse, the pre-synaptic cell will release a fraction *uX* of the neural transmitters into the synapse and this amount of transmitters will bind to the receptors to become *Y* and produce an EPSC in the post-synaptic cell (Clements et al., 1992; Bredt and Nicoll, 2003). Physiologically, *τ*_*d*_ represents the uptake process of the active transmitters back into the pre-synaptic cell to become *Z*, while *τ*_*r*_ represents the repackaging process of the neural transmitters so that they can be in the ready-to-release pool, *X*, again (Rizzoli and Betz, 2005; Alabi and Tsien, 2012).

In the TUM picture, the clearance of the glutamate from the synapse is carried out through the uptake by the pre-synaptic cell. However, it is known that fast glutamate uptakes by glutamate transporters (GluTs) are carried out both at pre-synapse cell and mostly through astrocytic processes to ensure a rapid clearance and thus decay of the EPSCs (Danbolt, 2001). In order to include the effects of astrocytes on the clearance process, we introduced a new pathway as well as a new state to the TUM model allowing the uptake of the synaptic glutamate by the astrocytes. This is referred to as the TUMA model as described in the Figure 7A.

The state *A* is the fraction of glutamate in the astrocytes. From the diagram, *A* comes from *Y* in the synapse and can be transformed into *Z* and then finally to *X*. The time scales *τ*_au_, *τ*_nu_, and *τ*_*g*_ are for the astrocytic uptake, the neuronal uptake, and the inactivation in the *A* state respectively. In this model, the released glutamate (*Y*) are transformed back into *Z* through two modes: a fast and a slow recycle pathways. The fast path takes place through direct pre-synaptic neuronal uptake of glutamate from the cleft within a timescale of *τ*_nu_ by EAAT3 (EAAC1) group of GluTs (Danbolt, 2001). For the slow path, glutamate in the synapse are first uptaken by the astrocytes via EAAT2 (GLT-1) group of GluTs with a time scale of *τ*_au_. These uptaken glutamate are then converted into neutral molecules, glutamine (Bröer and Brookes, 2001; Waniewski and Martin, 1986) and then converted back to the glutamate (Bak et al., 2006) once they are transported back into the pre-synaptic cell (*Z*). This whole process of glutamate conversion, release, uptake, and re-conversion to glutamate is slow and represented by a long time scale of inactivation *τ*_*g*_. Therefore, one can expect *τ*_*g*_ ≫ *τ*_nu_ and *τ*_au_. These ‘recovered’ glutamate are now readily loaded into the recycled vesicles at the pre-synapses and retrieved into active zone within *τ*_*r*_. The mechanism described thus takes the following form of equations:

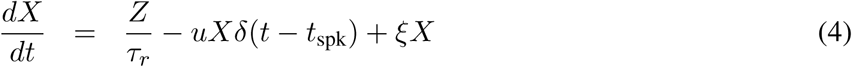

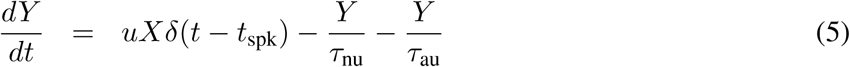

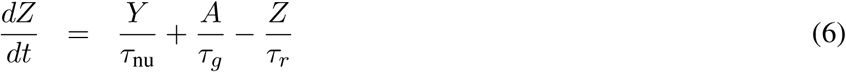

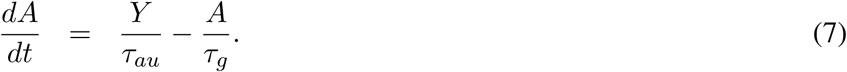

Note that the term *ξX* is added to the Eq. 4 because it is known that glutamate can be released either spontaneously or through the trigger of an action potential. The noise term for the asynchronous release as an extension to the TUM model was first studied by Volman et al. (2007). The goals of Volman’s work was to understand the dynamics of reverberation in neuronal cultures through reproduction of the experimental observations in the model with calcium dynamics. The noise term added to Eq. 4 was identified as spontaneous release events driven by the pre-synaptic residual calcium dynamics. Similar to Volman et al. (2007), the calcium-dependent asynchronous release was found to be essential for the model to reproduce the experimental observations. Therefore, we adopted their mechanism for the construction of TUMA model.

### Simulation

#### Neuronal Model

Similar to Volman et al. (2007), we implemented Morris Lecar (ML) neurons (Morris and Lecar, 1981) to test our synaptic model. The dynamics of ML neurons are described by,

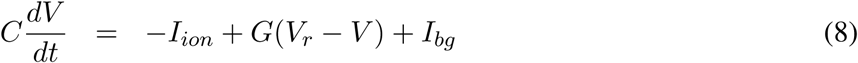

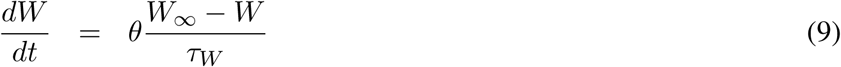

where

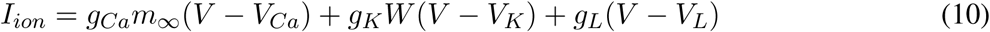

is the current through membrane ion channel. The variables

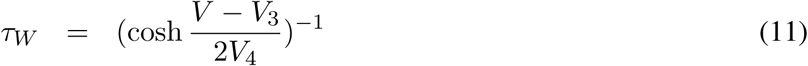

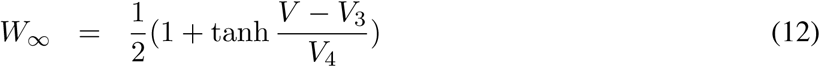

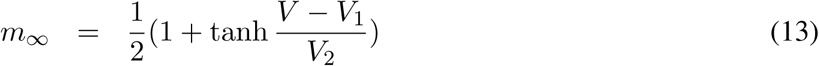

are the voltage dependent dynamic parameters. Another threshold parameter *V*_th_ of membrane potential is used to define the spiking events resulting in synchronous releases of neural transmitters at efferent synapses.

Additionally, a residual calcium variable *R*_Ca_ driven by the spiking events, as described by Volman et al. (2007), was also incorporated to produce reverberatory dynamics observed in cortical networks. The dynamics of residual calcium *R*_Ca_ is described by the equation,

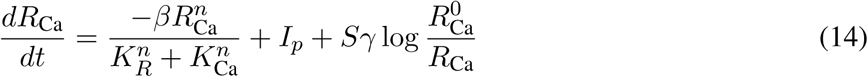

where *S* = Σ_*σ*_ *δ*(*t* − *t*_*σ*_) is the spike train with *t*_*σ*_ being the spike time. The residual calcium is used to determine the rate,

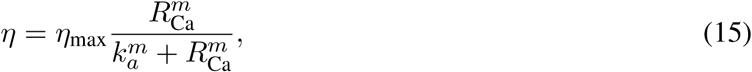

of synapse-dependent asynchronous releases of neural transmitters following an independent Poisson pro-cess at each efferent synapse. The neural transmitters released by the spike-driven synchronous and calcium-dependent asynchronous events follow a four state decaying dynamics based on the TUMA synapse description, Eqs. 4–7. In Eq. 4, the variable 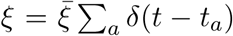 is then a summation of the asynchronous release events *a* with the Poisson rate given by Eq. 15. The parameters used are listed in the Table 1.

**Table 1:** Pharmacological targeting glutamate transporters alters SB statistics. Summary of the changes in array-wide neuronal activities from reference (Ref) after pharmacological treatments with glutamate transporter inhibitors; DL-TBOA(10*µ*M), DHK(200*µ*M) or selective augmentation of GLT-1 glutamate transporter with PAM (positive allosteric modulator) drug GT949(10*µ*M). Statistical tests: paired *t* tests, significance with Ref as baseline was defined as *p*<*0.05, n denotes number of cultures.

| Drug treatments | Firing rate (spikes/s) | SB rate (minute<sup>-1</sup>) | SB duration (s) | SB index (0-1) |
| --- | --- | --- | --- | --- |
| 1. Ref<br>+ DL-TBOA | 25±13<br>50±30, **<br>n=13 | 6±5<br>5±4, n.s<br>n=16 | 0.28±0.09<br>0.72±0.44, ***<br>n=16 | 0.75±0.12<br>0.84±0.11, *<br>n=13 |
| 2. Ref<br>+ DHK | 26±8<br>46±13, ***<br>n=11 | 6±5<br>6±6, n.s<br>n=11 | 0.34±0.17<br>0.66±0.39, **<br>n=11 | 0.84±0.12<br>0.67±0.18, **<br>n=10 |
| 3. Ref<br>+ GT949 | 29±14<br>15±10, ***<br>n=14 | 5±2<br>4±2, n.s<br>n=11 | 0.27±0.12<br>0.18±0.06, *<br>n=9 | 0.80±0.16<br>0.83±0.09, n.s<br>n=12 |

#### Network topology and connectivity

A network is constructed with 100 Morris–Lecar neurons randomly connected (connection probability = 0.1) with each other through the TUMA synapses. The inhibitory-to-excitatory ratio for the neurons is set to 0.2 (Soriano et al., 2008). The fractions of neurotransmitters in the active state *Y* multiplied by the synaptic weights 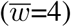, a scale for density of post-synaptic effectors such as glutamate receptors, determine the contributions of the afferent synapses to the membrane conductance *G* of the post-synaptic neurons,

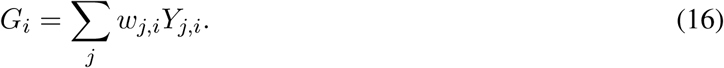

Here, we assume a linear summation over all pre-synaptic neurons *j* for the given post-synaptic neuron *i*. The synaptic weights *w* were randomly drawn from a Gaussian distribution truncated at its width that is set to *±*20% of its mean 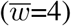 for the connected neurons.

#### Implementation and Code Accessibility

A GUI program, written in C++ programming language using the Common Simulation Tools framework in previous work of Huang et al. (2017a) implementing the Volman’s model was used to perform the simulations. The program was modified as per our TUMA synaptic dynamics for the s imulations. Original package measim implementing Volman’s synaptic model is available on GitHub (https://github.com/chnchg/measim). Spike data obtained from the simulation were exported to the MATLAB software for further processing and analysis. The same algorithm for SB detection (described above) was used for comparison with experimental results.

### Statistical Analysis

Assumptions of normality were validated using D’Agostino-Pearson omnibus test (*α* = 0.05). Hypothesis testings were performed on repeated measures using parametric one-way ANOVA or two-tailed paired/unpaired *t* tests. In case of one-way ANOVA in multiple groups, Tukey’s post-hoc multiple-comparison tests were also performed. Non-parametric Friedman test was performed for multiple group comparison in case any one group failed the normality test. Sample sizes are listed in the figure captions. Error bars indicate standard error of the mean (SEM) unless otherwise specified.

## Results

### SB associated active and dormant states in cortical networks

Neurons in cortical cell cultures develop into self-organized networks forming recurrent connections (Figure 1A). At the same time, astrocytes also develop and extend their processes forming their own complex network, at the vicinity of neuronal network. Functionally, the neurons exhibit a wide range of spontaneous firing patterns (Wagenaar et al., 2006) during their course of development. Typically, after three weeks of age neuronal connections become functionally mature, and the array-wide firing patterns turn stable. Each electrode of MEA generally display two patterns of neuronal activities: either bursts of spikes followed by isolated spikes (ch#2,3,7,9,10,14 in Figure 2A), or only bursts of spikes (ch#1,4,5,6,8,11,12,13). Aligning the temporal activities in all the electrodes, the overall array-wide firing dynamics thus appears as either asynchronous or synchronous firings in a network. A distinction between synchronous bursting and asynchronous spiking events can be made by generating inter-spike interval (ISI, time interval between successive discharge) return map (Figure 2B). Electrodes which exhibit only synchronous events, their ISI return map show single cluster lying within the range of 2–250 ms. Whereas, electrodes which also show asynchronous firings, display additional clusters of ISI distributed in the interval between 1–10 sec.

**Figure 1:**
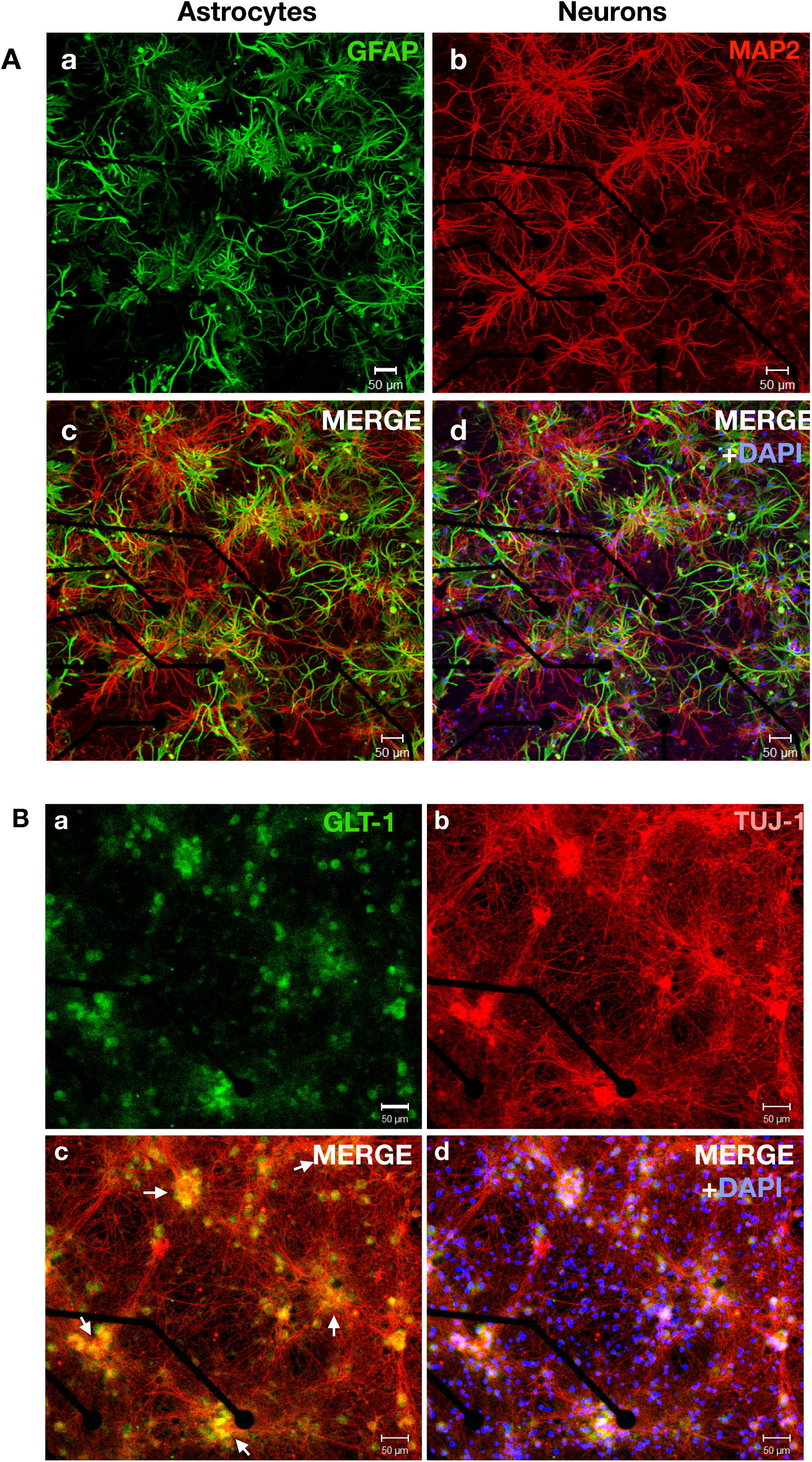
Astrocytes and neurons develop into complex networks in cortical cultures. (A) Confocal micrograph of a 22 DIV old cortical network fixed on multi-electrode arrays. (A-a) GFAP immuno-stained (green) cortical network shows highly complex dense structures of stellate type astrocytes coexisting with highly interconnected network of MAP2-stained (red) neurons (A-b). (A-c) Merged image of GFAP and MAP2 staining, shows tightly interlinked neuron-astrocyte network. (A-d) shows co-localized image with along with DAPI signals. (B) Confocal micrograph of a 14 DIV old fixed cortical network fixed on a sample multi-electrode array. Panel (B-a) shows immuno-stained signals of GLT-1 (green) glutamate transporters distributed as clusters. (B-b) Neurons were stained for TuJ1 (red), a marker for neuronal cytoskeleton. Neurons can be seen to be randomly organized as thick clusters with hub-like topology. (B-c) Merged GLT-1 and Tuj-1 staining signals show that GLT-1 were strongly expressed and localized at the hub locations where neuronal cell bodies form dense synaptic clusters (arrows). (B-d) shows co-localized image with along with DAPI signals.

**Figure 2:**
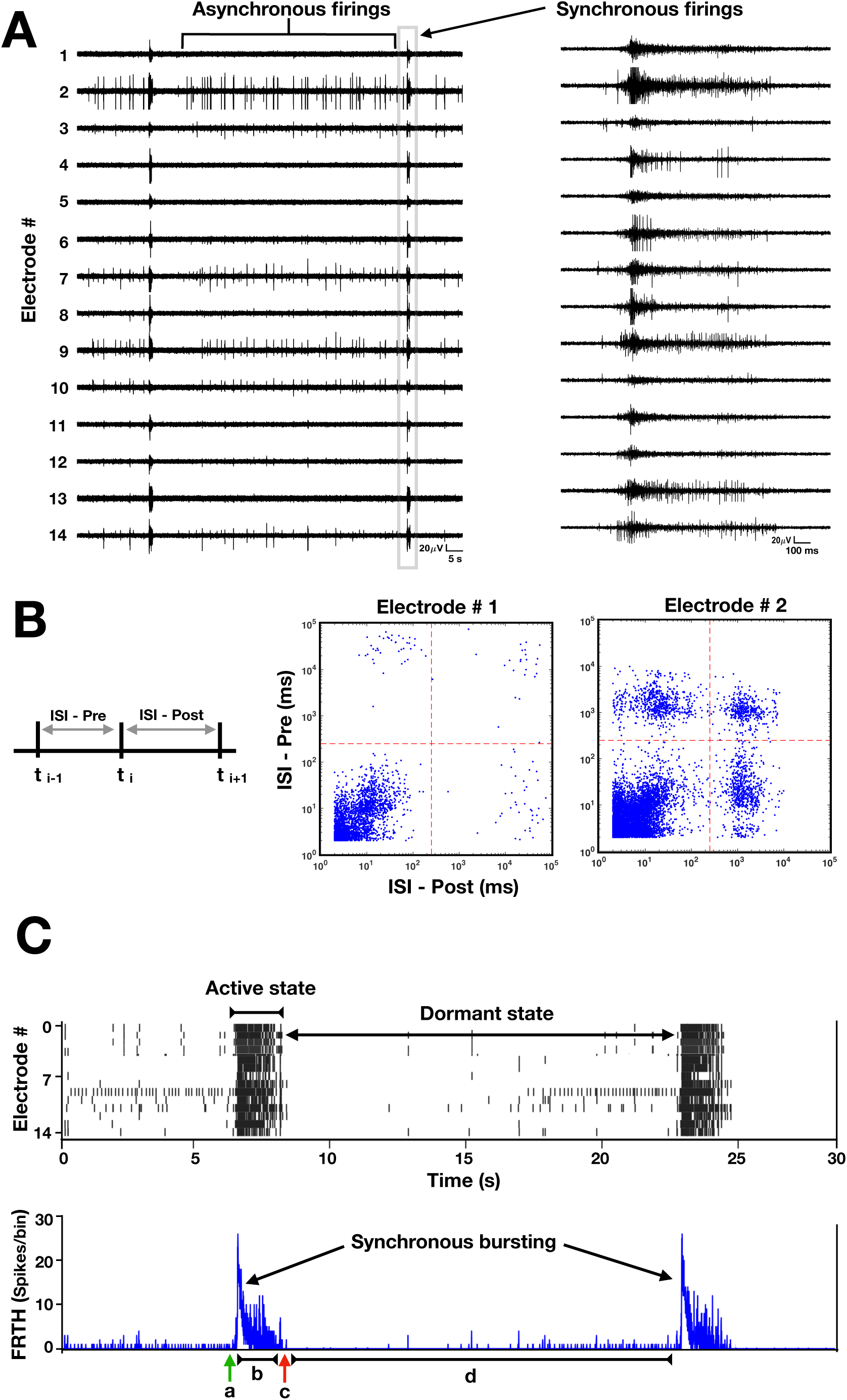
Active and dormant states of neurons in cortical cultures. (A) Time series of extracellular field potentials of a sample cortical network, recorded with 14 electrodes using a multi-electrode array probe. Spontaneous electrical activities of cortical neurons in a developing network (22 days *in vitro*) appear as asynchronous or synchronous firings. (B) A inter-spike interval (ISI) distribution, generated by plotting time intervals (top) between successive firings, show a distinction between synchronous and asynchronous firing events. Electrode#1 engaging only in synchronous firings exhibit only one dense cluster of ISI lying in 2–100 ms range (middle), while the electrode#2 also engaging in asynchronous firings show additional ISI clusters lying above 1 sec region (bottom). (C) Raster plot of detected spikes (top, black trace) illustrates two states of network in general: ‘active’ state when all units exhibit synchronous burst (SB) firings followed by a ‘dormant’ state in which most units show no to very few activities. Corresponding firing-rate time histogram (FRTH) (bottom trace in blue), calculated by summing all the detected spikes within 5 ms bin size, displays the kinetic structure of network’s states. The kinetics involve an initiation (a), maintenance (b), and termination (c) phases followed by a much longer dormant phase (d).

We will refer to the synchronous-discharge state of network as the ‘active state’, since all neurons collectively remain active during this state (Figure 2C). Within an active state a network exhibits different kinetics, as shown in the firing-rate time-histogram (FRTH) plot. FRTH was generated by summing the total number of array-wide action potentials detected in a 5 ms bin size. A generic SB’s kinetic structure consists of the initiation, maintenance, and termination phases (Figure 2C, bottom). Initiation of SB occurs by assembly of neuronal activities at an exponential rate (Eytan and Marom, 2006). Post-assembly of neuronal firings, array-wide firing reaches a peak level then gradually decays to the extent where all neuronal spikes ceased completely. By taking reciprocal of the ISI ranges stated above, it can be seen that the firing rate in single electrode during the active state can range between 10-500Hz. After termination of the SB, the network maintains a period of quiescence for several seconds and regains its basal firing activities until next SB occurs. We will refer to the time between the termination and the initiation of the next SB as the ‘dormant state’ of the network. During the dormant state some neurons may still exhibit spontaneous firings at a rate within the range of 0.1-4Hz.

### Synaptic glutamate dynamics accompanying SB

Firing dynamics observed in SB involves fast recurrent excitation of AMPA and NMDA receptors by synapse-released glutamate (Clements et al., 1992; Bredt and Nicoll, 2003). Previous studies indicate that most of the synaptic glutamate(≈80%) is uptaken by astrocytic processes through GLT-1 transporters while only a smaller portion is directly uptaken by the neurons (Lehre and Danbolt, 1998; Eulenburg and Gomeza, 2010). This distinction is due to the difference in the number of transporter expressions on the membranes of the two cell types. We performed a qualitative check on GLT-1 expression in our sample cortical cultures. Confocal imaging, revealed a highly clustered expression of GLT-1 (see Figure 1B) around hub-like clustered neuronal bodies. Kavalali et al. (1999) has already shown these neuronal hub structures to be composed of organized synaptic clusters formed by assembly of presynaptic, postsynaptic and dendritic structures. Observation of intense GLT-1 expressions implies high glutamate activities at these locations. Our goal was to detect glutamate dynamics during SBs in the immediate surrounding of the astrocytes. Therefore, we employed glutamate sensors genetically expressed on astrocytic-membrane facing extracellular space.

Astrocytes in developing cortical cultures were infected with adeno-associated viruses expressing GFAP-targeted intensity-based-glutamate-sensing-fluorescent-reporter (iGluSnFr) (Marvin et al., 2013) (See material and methods for details). This approach has been previously shown to indicate the kinetics of glutamate clearance by astrocytes (Armbruster et al., 2016). Therefore, we implemented this method for evaluating GluT activities during SBs. Within two weeks (post-infection), fluorescence signals from astrocytes were detectable. Simultaneous recording of MEA and whole field synaptic glutamate imaging, revealed sharply elevated iGluSnFr signals associated with SB events recorded with MEA(Figure 3A). Its kinetics almost overlapped with the time course of network’s active state. Evidently, the sharp rise in glutamate signal was due to synaptic releases of glutamate during intense burst firings by neurons. Notably, the glutamate level rapidly decayed back to baseline level immediately towards the termination of the SBs(Figure 3B). Detected iGluSnFr signal continued to remain at its baseline level until the next SB occurred. The time taken for the decrease of glutamate from its maximum level to half during SB was defined as *τ*_half_. The glutamate decay time *τ*_decay_ (assuming exponential decay function (Armbruster et al., 2016)) was computed as *τ*_half_*/* ln 2 (Figure 3C) to reflect the glutamate clearance time by the astrocytes. In our culture system, the decay time was found to be distributed around 0.26*±*0.11 s (N=8 cultures).

**Figure 3:**
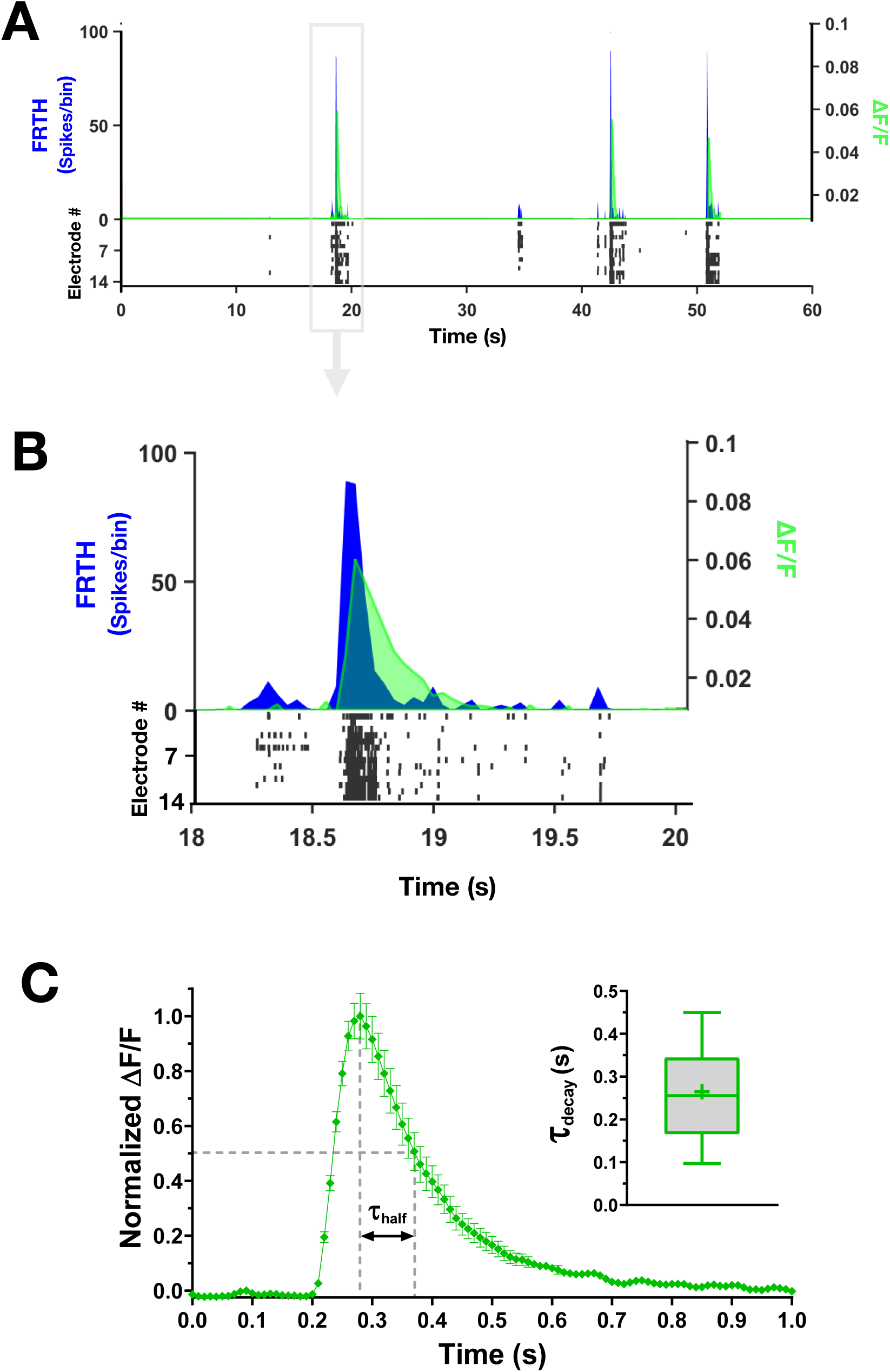
Synaptic glutamate level rises and falls during SBs. Simultaneous recording of neuronal activities with MEA and synaptic glutamate dynamics detected by astrocytes with iGluSnFr imaging. (A) Top panel shows summed neuronal spiking activities as firing-rate time histogram (FRTH with bin size 40ms, blue trace) overlapped with glutamate dynamics detected simultaneous through iGluSnFr imaging. Change in fluorescence is plot as Δ*F/F* (green trace). Bottom raster plot (black) shows detected neuronal activities from individual electrodes. (B) shows enlarged view of one of the SB events in A. Note that synaptic glutamate level increases globally and decays rapidly during and after SB events. (C) Average normalized fluorescence Δ*F/F* from a sample network (green trace). The time taken for decrease of synaptic glutamate level from its maximum to half during SB is denoted by *τ*_half_. Inset, summary of glutamate decay time *τ*_decay_ in cultured cortical networks, calculated as *τ*_half_*/* ln 2. Middle line in the box plot shows median and cross represents mean(0.26*±*0.11 s, n=8 cultures).

### Pharmacologically targeting astrocytic GLT-1 alters SB properties

To gain further insights on the role of glutamate transporters in SBs, we used pharmacological tools to dissect their contribution. Adding GluT-specific inhibitor drugs resulted in dramatic change in firing and bursting patterns (Figure 4B-C). Inhibition with DL-TBOA resulted in two-folds increase in array-wide firings (Table 1). Still there was no significant change in SB frequency. The SB duration was increased nearly 2.5 times. The level of array-wide spiking activities during SBs was quantified as SB index (ranging between 0 and 1, see material and methods for more information). Our cultured networks usually displayed an SB index of ≈0.8 in their reference recordings. DL-TBOA treatment significantly increased SB index. We also observed a rearrangement effect of firing activities in different samples. Although all networks showed consistent increase of firing activities after DL-TBOA addition, a range of change in SB duration was observed across samples. Networks which showed small increases in burst duration after the drug treatment, showed increased frequency of SB compared to their reference. Whereas, networks that displayed big increase in burst duration, showed a decrease in SB frequency. Specific inhibition of GLT-1 with DHK reproduced the effects similar to DL-TBOA treatment. The only difference was found in the SB index of the networks after drug treatment. Contrary to DL-TBOA effect, DHK addition resulted in increased asynchronous state firings which significantly reduced the burst index (Table 1). On the other hand, augmenting GLT-1 function with GT949, a positive allosteric modulator (PAM) drug, significantly decreased array-wide firings without affecting SB frequency and SB index. The duration of SBs became shorter than their reference, a consequence likely due to reduced firing activities (Figure 4D). These results clearly show that the synaptic efficacy during SBs is regulated by GLT-1.

**Figure 4:**
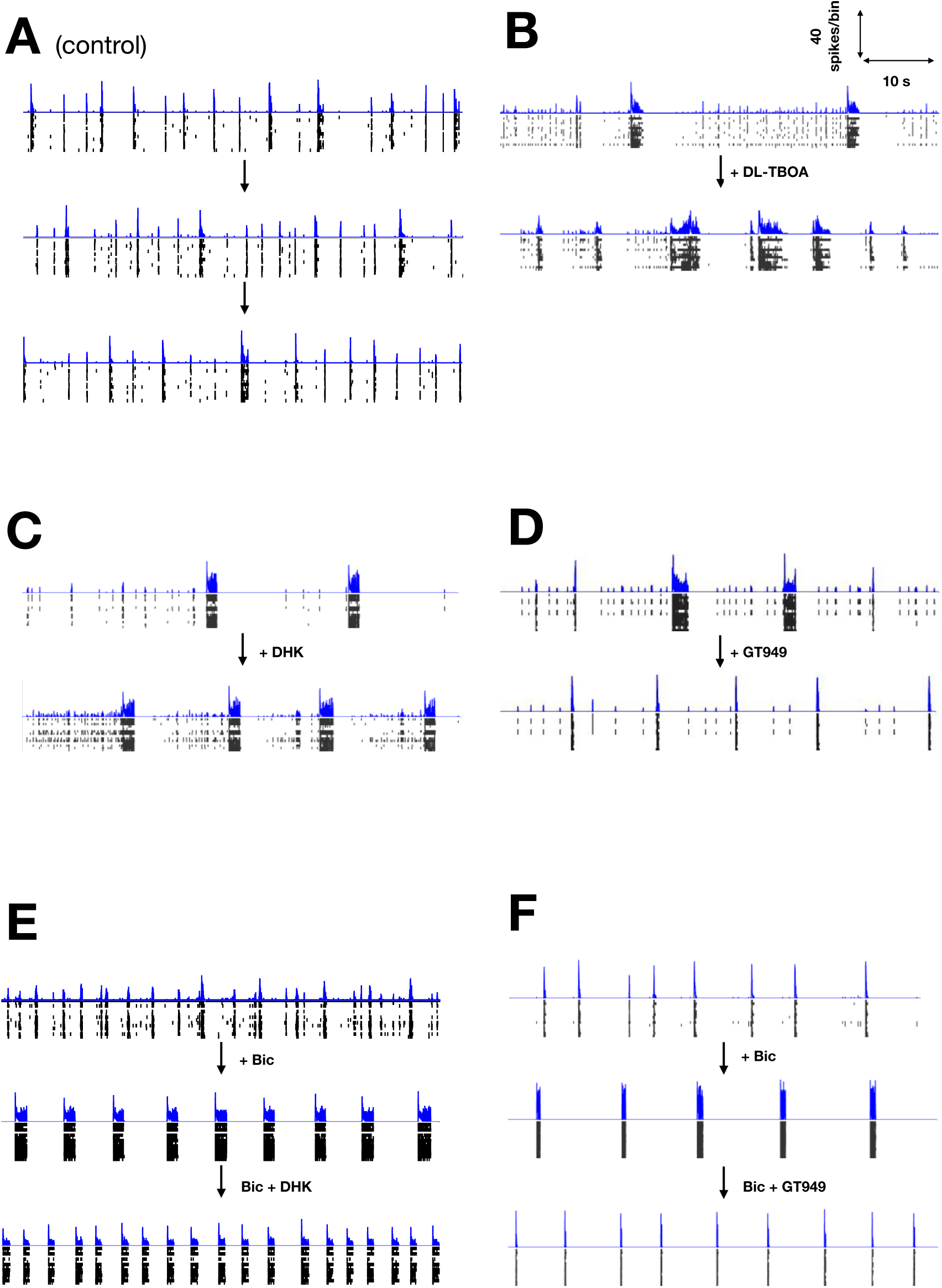
Pharmacological manipulation of glutamate transporters alters SB dynamics. Figures(A–F) shows one-minute snapshots of firing activities obtained from six different cortical networks. FRTHs generated with a bin size of 5ms are shown in blue while the corresponding raster plot of detected spikes are shown in black. The interval between each snapshot of a network, shown in the order as indicated by the arrow marks, were at least separated by 45 minutes. The first snapshot of each network shows its reference activities level. Network in A shows the time effect on network dynamics in a long duration recording. Without any pharmacological treatment network maintained similar firing and bursting patterns over a period of 120 minutes. Network in B, after treatment with DL-TBOA (10*µ*M), resulted in prolonged and frequent SBs. The dormant states were shorter and contained with lesser asynchronous activities. Similar effects were observed after DHK (GLT-1 specific, 200*µ*M) treatment in network in C. The only difference between DHK and DL-TBOA was an increase of asynchronous activities during the dormant state under DHK. Specific augmentation of GLT-1 (10*µ*M) of network in D with GT949 resulted in shortening of SBs. GLT-1’s role was further tested in disinhibited state of cortical networks. Disinhibition with bicuculline (Bic, GABA inhibitor, 10*µ*M), shown in network in E, resulted in more robust SB patterns with clear active and dormant states of network. Resulting SBs were longer than its reference. Further DHK (200*µ*M) addition, enhanced SB frequency and reduced SB duration at the same time. Enhancing GLT-1 function with GT949 (10*µ*M) treatment in disinhibited state (network in F), resulted in shorter SBs.

### SBs are not terminated by inhibition

Inputs from the inhibitory sub-networks are known to regulate the excitability of neurons. It is the interaction between the two which is believed to determine the flow of information through the network as well as shape network activities (Ziburkus et al., 2006; Kudela et al., 2003; Chen et al., 2006; Iida et al., 2018). To gain insight on the contribution of inhibitory inputs in SB, cortical networks were treated with bicuculline (Bic, GABA antagonist, 10 *µ*M) to suppress the fast inhibitory effects of *γ*-aminobutyric acid (GABA). After bicuculline addition, the SBs became more periodic and longer(Figure 4E-F). Networks in disinhibited state also often displayed reverberatory activities (sub-bursts) within each SB event similar to previous report of Lau and Bi (2005). Asynchronous activities were almost abolished while SBs were more prominent, with rather clearly-maintained active and dormant phases. The observation that networks did not fire endlessly and stop until there is no more activity after bicuculline treatment, shows that the mechanism of SB termination is not limited to excitatory-inhibitory interactions.

Analyzing the firing patterns, cultures treated with bicuculline showed significant increase in firing rate (Table 2). After disinhibition, networks engaged only in synchronous activities (burst index ≈ 1). Disinhibition also resulted in significant increase in the duration of SB while maintaining similar SB frequency compared to reference. To validate the role of GLT-1 in mediating SB dynamics in disinhibited state of cortical networks, we further pharmacologically manipulated GLT-1 transporters in the the disinhibited state of network. Further addition of DHK did not affect the overall firing rate or the SB index but induced a re-organization of the firing events into shorter duration with higher frequency of bursting (Figure 4E). These effects were reversible after washout of the drugs (data not shown). Adding GT949 in the disinhibited networks significantly decreased the overall network firing activities, while maintaining the same high SB index induced from disinhibition (Figure 4F). Under the influence of GT949, burst duration was significantly shorter, without affecting the SB frequency (Table. 2).

**Table 2:** Comparison of SB statistics under GLT-1 specific treatments in disinhibited state of cortical networks. Summary of the changes in array-wide neuronal activities from reference (Ref) to pharmacologically disinhibited state (Bic, 10*µ*M) followed by manipulation of GLT-1 glutamate transporters with DHK (200 *µ*M) or GT949 (10 *µ*M). Statistical tests: repeated measures one-way ANOVA (F) and non-parametric Friedman test (*χ*^2^), significance *p*<*0.05.

| Drug treatments | Firing rate (spikes/s) | SB rate (minute<sup>-1</sup>) | SB duration (s) | SB index (0-1) |
| --- | --- | --- | --- | --- |
| 1. Ref | 34±22 | 7±5 | 0.28±0.19 | 0.86±0.12 |
| + Bic | 82±48, * | 7±4, n.s | 0.85±0.49, * | 0.99±0.01, * |
| + DHK | 97±31, n.s | 21±16, * | 0.58±0.35, * | 0.99±0.01, n.s |
| | F(2,10)=21.19, *** | F(2,10)=10.26, ** | F(2,8)=12.21, ** | $\chi^2$ (2,9)=9.8, ** |
| 2. Ref | 27±17 | 5±2 | 0.35±0.24 | 0.84±0.12 |
| + Bic | 60±42, * | 8±4, n.s | 0.73±0.42, *** | 0.99±0.01, ** |
| + GT949 | 12±08, ** | 5±3, n.s | 0.17±0.05, ** | 0.99±0.02, n.s |
| | F(2,9)=14.88, ** | F(2,8)=2.43, n.s | F(2,9)=16.64, ** | $\chi^2$ (2,8)=12.67, *** |

### Synchronous calcium elevation in astrocytes is induced by SBs

Previous studies have reported astrocytic calcium sensitivity towards synaptic glutamate activities (Dani et al., 1992). However, its role in synaptic events still remains unclear. We further probed calcium activities in astrocytes in spontaneously bursting cortical networks. Cultures were expressed with genetically encoded calcium sensors (cyto-GCaMP6f) specific to the astrocytes. GCaMP6f imaging revealed asynchronous local calcium elevation as well as synchronous calcium elevations (SCEs, Figure 5A). We noticed that, the sequence of astrocytic activation during the global calcium elevation was not random. A hierarchy in the recruitment of calcium was consistently observed. This was likely due to an existence of a stable subset of privileged neurons whose spiking activities reliably increase before the onset of global SB (Eytan and Marom, 2006). Combining MEA and calcium imaging, temporal activities from the two cell types were recorded simultaneously. Consistently, we observed that SCEs always emerged towards the end of longer persisting SBs (Figure 5B). Note that during the SCE in astrocytes, neuronal ongoing activities were completely abolished. Neurons resumed their basal activity level shortly after decay of SCE in astrocytes. Importantly, only SBs persisting long enough (*>*500 ms) were followed by the astrocytic SCE. Since SBs result in large glutamate transient at synapses globally, this transient is likely to triggered collective activation of astrocytic metabotropic glutamate subtype 5 receptors (Panatier et al., 2011). A positive correlation was indeed found between the duration of SB in neurons and its concomitant SCE in astrocytes. To validate whether glutamate released by the SBs were responsible for SCE in astrocytes, cultures were treated with riluzole (glutamate release inhibitor, 10 *µ*M). Treatment with riluzole resulted in suppressed neuronal activities with complete abolition of SBs and associated SCEs (data not shown). Although neurons maintained some asynchronous firings, no astrocytic SCE was observed after addition of riluzole, indicating SB dependency of synchronous calcium activities in astrocytes.

**Figure 5:**
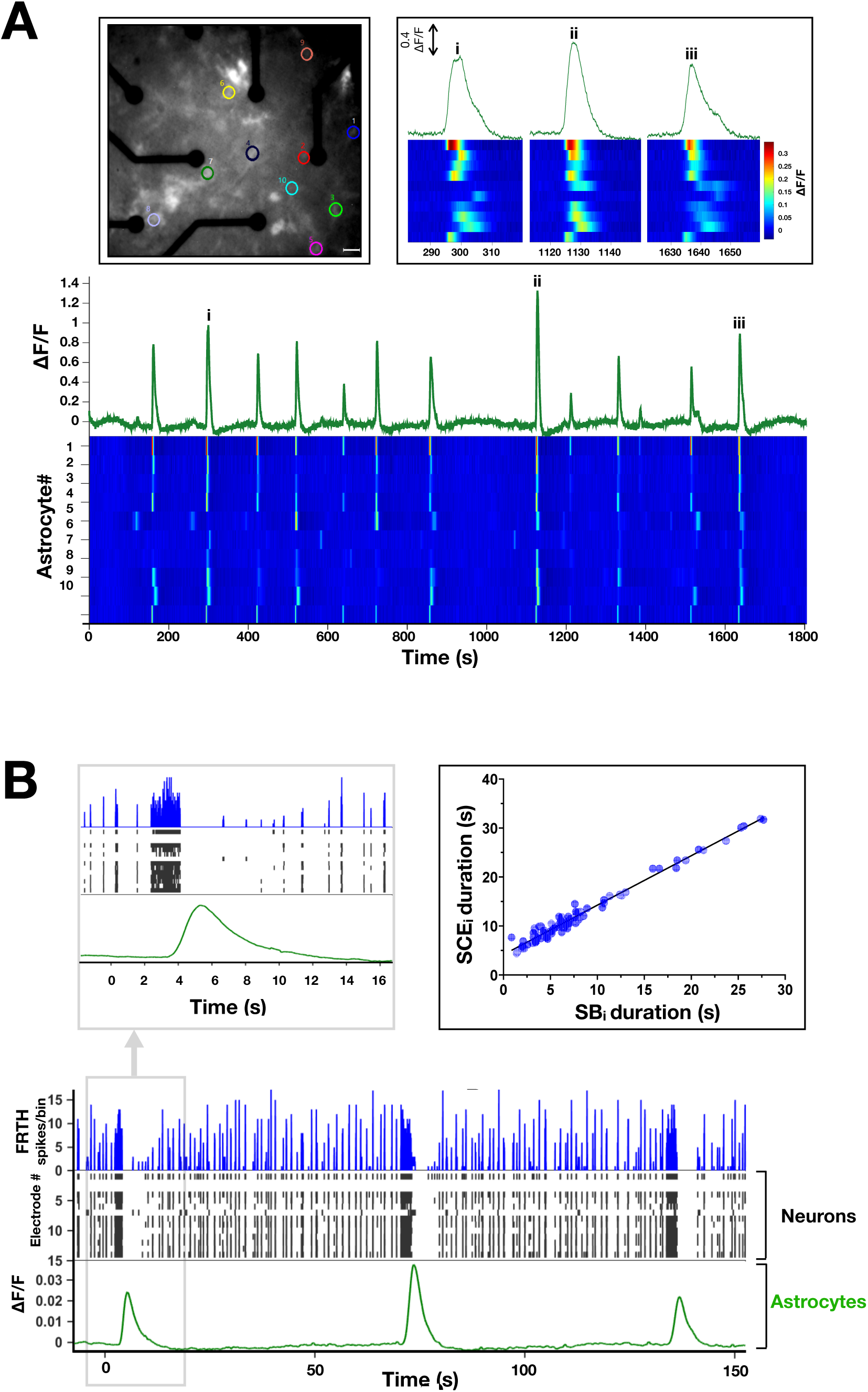
Astrocytes exhibit synchronous calcium dynamics following SBs. (A) Top inset on left shows fluorescence micro-graph of a culture expressed with cyto-GCaMP6f calcium sensors in astrocytes, scale bar – 30 *µ*m. Different colored circles indicate locations of 10 randomly selected and spatially distant calcium-active astrocytes. Traces in green show summed fluorescence intensity–time plot of the selected astrocytes. Heatmap shows corresponding temporal dynamics of calcium intensity in individual astrocytes. Top right inset shows enlarged view of three synchronous calcium elevation (SCE) events marked as (i,ii,ii). Note that the activation pattern remains similar in each event, indicating a hierarchical order of astrocyte activation during SCEs. (B) Shows temporally aligned activities of neurons obtained from MEA (FRTH with 40 ms bin size, blue trace) and corresponding raster plot(black trace), with collective calcium activities in astrocytes. Top left inset, shows enlarged view of a global event when an astrocytic SCE appears when a long persisting neuronal SB occurs. Note that all neuronal activities ceased during the global calcium rise in astrocytes. Neuronal activity reappeared soon after the SCE decay. (Top right inset) Duration of SCE was found to be linearly related to its preceding SB duration(*n* = 87, 4 cultures).

### Simulation results

The description of firing and bursting patterns obtained from MEA experiments show how the dynamics of SB changes under various conditions. We used these firing patterns as input to obtain relevant parameters for our tripartite synapse TUMA model. This model describes how glutamate gets relocated to different parts of a tripartite synapse. For model details, please refer to the material and methods. The essentials of this model can be understood from the schematic diagram shown in Figure 6A. The amount of glutamate (expressed as fractions) in the system can be in *X* (ready-to-release pool), *Y* (active state; producing EPSC), *Z* (pool of glutamate uptaken from the synapse), and *A* (glutamate uptaken by astrocyte). Similar to the TUM model (Tsodyks et al., 2000), we implemented conservation of glutamate in the tripartite synapse; with *X* + *Y* + *Z* + *A* = 1. The glutamate transporters targeted in this study are related to the two uptake time scales in TUMA synapse; *τ*_nu_ (neuronal uptake) and *τ*_au_ (astrocytic uptake). The values of *τ*_nu_ and *τ*_au_ were obtained and fixed in a manner to match the simulated firing patterns with the experimental observations. These time constants and the model mechanism were the basis of our understanding on how glutamate trafficking dynamics regulate the SB properties.

**Figure 6:**
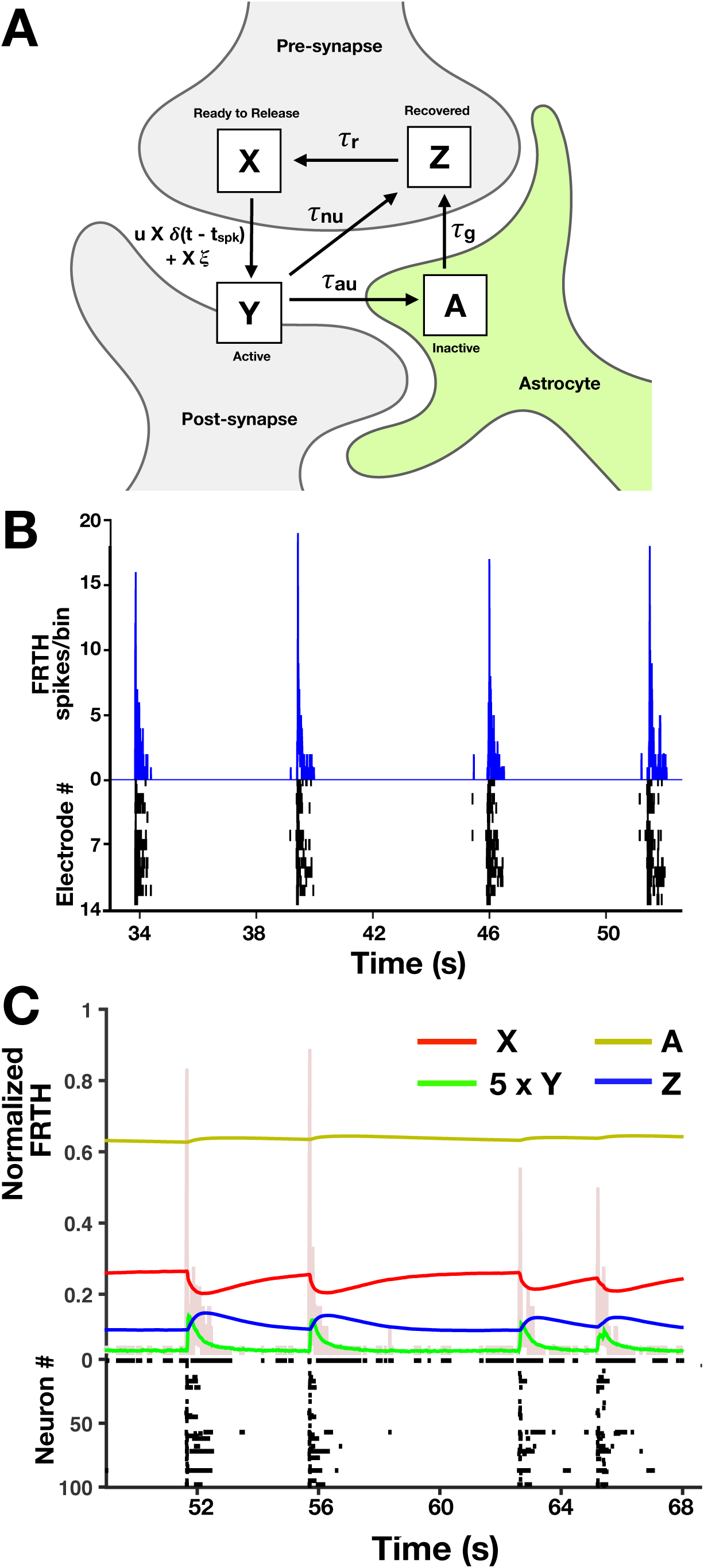
Simulation with TUMA synapses reproduces experimental SB patterns. (A)Schematic representation of TUMA, tripartite synapse model describing glutamate recycling. The fraction of the total amount of the glutamate in these various states are represented by *X* (in presynapse), *Y* (at synapse), *Z*(recovered at presynapse) and *A*(in astrocytes) as shown in the figure and *X* + *Y* + *Z* + *A* = 1. Impulses *δ*(*t* − *t*_spk_) created by spikes at time *t* = *t*_spk_, arriving at the presynapse, trigger glutamate release into the cleft. Released glutamate molecules rapidly activate the post synaptic fast receptors until their clearance by neuronal and astrocytic uptake with timescales *τ*_nu_ and *τ*_au_ respectively. The astrocytic uptake follows glutamate–glutamine cycle leading to slow recovery of glutamate at the timescale of *τ*_*g*_ in the presynapse. Once recovered, glutamate gets packed into vesicles and retrieved back at the active zone within *τ*_*r*_. During this recovery process, some neurons may exhibit noise(*ξ*) induced asynchronous release. (B) Shows FRTH (blue trace, bin size 5ms) and raster plot (black trace) of neuronal activities obtained from MEA recording. (C) Similar firing and bursting patterns were obtained from simulation (random network of 100 neurons connected with TUMA synapses). Simulated normalized FRTH trace (bin size 5ms) is overlapped with the four-state *X*(red), *Y* (green), *Z*(blue), and *A*(yellow) dynamics of glutamate occurring simultaneously in the network. These traces denotes the average fraction of glutamate distributed around the synapses. During SBs, a very small fraction of the total glutamate gets released. Therefore, for visual clarity, *Y* is shown 5 times its actual values.

### Synchronous bursting events

Figure 6C shows the raster plot of typical noise-induced synchronous firing events in 100 neurons randomly connected through TUMA synapses. The time course of *X, Y, Z*, and *A* state of glutamate averaged across all synapses during the SB events are also shown in the figure. Simulation parameters (see Table 3) were chosen such that the firing patterns were similar to experimental firing patterns (Figure 6B) and consistent with previous reports (Volman et al., 2007; Huang et al., 2017a). Prior to the onset of SB, the fraction *X* remains at its maximum, while *Z* at its minimum. Most of the changes in *X* gets supplied from *Z*. Once the SB is triggered, a steep rise in *Y* happens similar to the experimental observations (Figure 2). One remarkable feature shown in the figure is that during an SB, changes in *X* and *Z* are large while only very small changes occurs in *A*. These changes in *A* during SB are small because of the slow process for *A* to turn into *Z*. If this later process could be faster, one would expect bigger changes in *A* during an SB event.

**Table 3:** List of parameters used in simulation. With the exceptions of *τ*_au_ and *τ*_*g*_, all the other parameter values used in this work were obtained from previous works (Tsodyks et al., 2000; Volman et al., 2007; Huang et al., 2017a). Through out all the simulation runs, the leaking channel conductance parameter(*g*_*L*_) was set slightly lower (0.46mS) than the previous works (0.5mS) to induce more spiking activities. The value of release constant for residual calcium (*γ*) was used as 33 *µ*M s^-1^ (Huang et al., 2017a), whereas Volman et al. (2007) used 80 *µ*M s^-1^. In our simulations *γ* was tuned to 50 *µ*M s^-1^. After fixing the astrocytic uptake time constant *τ*_au_ equal to the experimentally obtained *τ*_decay_ value (see Figure 3C), glutamate recovery to the *Z* state by the neuronal uptake mechanism *τ*_nu_ was tuned to 50ms to match with the experimental firing and bursting patterns. Glutamate retrieval from the *Z* state to the ready to release active zone *X* was set (600ms) within the range of ‘kiss-n-run’ mode of vesicle retrieval *τ*_*r*_ (Gandhi and Stevens, 2003). All the other remaining parameters were identical to the previous works (Tsodyks et al., 2000; Volman et al., 2007; Huang et al., 2017a).

|  |  |  |  |  |  |
| --- | --- | --- | --- | --- | --- |
| $g_{Ca}$ | $1.1 \text{ mS.cm}^{-2}$ | $C$ | $1 \mu\text{F cm}^{-2}$ | $k_a$ | $0.1 \mu\text{M}$ |
| $g_K$ | $2 \text{ mS.cm}^{-2}$ | $\theta$ | $0.2 \text{ m s}^{-1}$ | $k_r$ | $0.4 \mu\text{M}$ |
| $g_L$ | $0.46 \text{ mS.cm}^{-2}$ | $I_{bgm}$ | $27 \mu\text{A}$ | $\beta$ | $5 \mu\text{M s}^{-1}$ |
| $V_{Ca}$ | $100 \text{ mV}$ | $I_{bgs}$ | $0 \mu\text{A}$ | $m$ | $4$ |
| $V_K$ | $-70 \text{ mV}$ | $u_{low}$ | $0.2$ | $n$ | $2$ |
| $V_L$ | $-65 \text{ mV}$ | $u_{up}$ | $0.2$ | $I_p$ | $0.11 \mu\text{M s}^{-1}$ |
| $V_1$ | $-1 \text{ mV}$ | $\tau_{nu}$ | $50 \text{ ms}$ | $\gamma$ | $50 \mu\text{M s}^{-1}$ |
| $V_2$ | $15 \text{ mV}$ | $\tau_r$ | $600 \text{ ms}$ | $[\text{Ca}^{2+}]_o$ | $2 \text{ mM}$ |
| $V_3$ | $0 \text{ mV}$ | $\tau_{au}$ | $250 \text{ ms}$ | $V_{th}$ | $10 \text{ mV}$ |
| $V_4$ | $30 \text{ mV}$ | $\tau_g$ | $30 \text{ s}$ | $w_{ee}$ | $4.0$ |
| $V_r$ | $0 \text{ mV}$ | $\xi_{ave}$ | $0.02$ | $w_{ei}$ | $4.0$ |
| $V_i$ | $-90 \text{ mV}$ | $\eta_{max}$ | $0.32 \text{ ms}^{-1}$ | | |

### Slower astrocytic uptake increases SB rate and duration

The effects of the astrocyte uptake time scale on the properties of SB are shown in Figure 7A(a–b). It can be seen that an increase in *τ*_au_, from 200 ms to 300 ms, resulted in a substantial increase in the overall firings, bursting rate, bursting duration and burst index (Figure 7B). This effect should be similar to the inhibition of the astrocytic uptake by DHK treatment in our experiments. These increases reflect that more glutamate becomes available in the synapse while *A* gets smaller. Figure 7A(a–b) shows that *A* remains relatively constant during the SBs however its mean value is sensitive to the value of *τ*_au_. A change of *τ*_au_ from 200 ms to 300 ms decreases *A* from 0.71 to 0.66. A further increase in *τ*_au_ would decrease *A* to an even smaller value and vice versa. These findings are consistent with the schematic picture that *A* works as a temporary storage for the glutamate and can regulate the amount of available glutamate in the synapse (Jabaudon et al., 1999). With the parameters used to reproduce experiment observations, the simulation revealed that *A* remains relatively constant and sets the amount of glutamate available to *X*.

**Figure 7:**
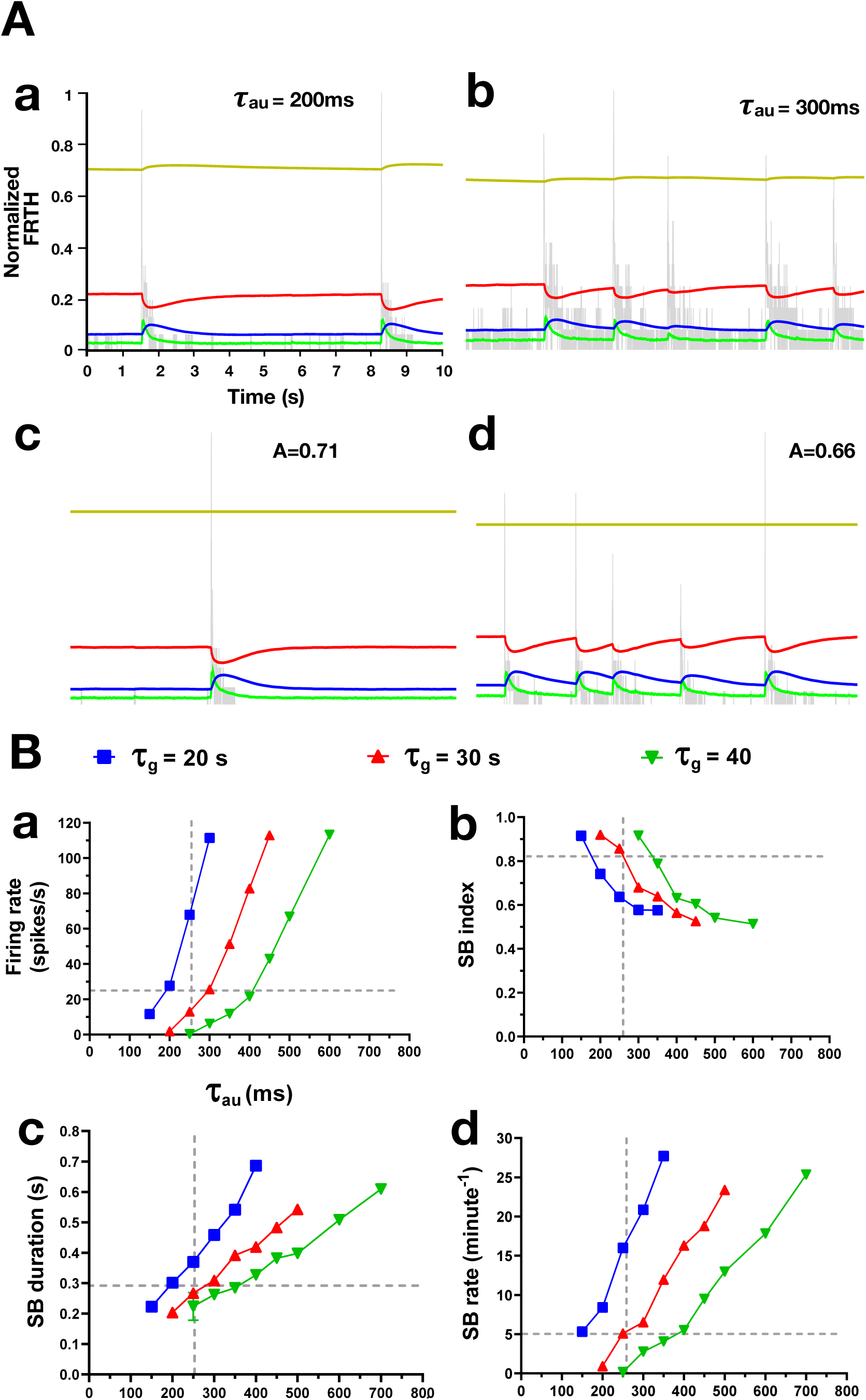
Astrocytes regulate the amount of glutamate available to neurons. (A) Effects of varying the astrocytic uptake timescale *τ*_au_ on network firing patterns (FRTH, gray trace). Subplots A(b–d) share the same vertical and horizontal axis scales with (A-a). A(a–b) Increasing *τ*_au_ from 200 ms to 300 ms resulted in longer and frequent SBs similar to DHK effects. A(c–d) By fixing *A* to the mean values (0.71 and 0.66) obtained from *τ*_au_ (200 and 300ms) did not affect the SB patterns. (B) Summary of network firing statistics dependence on astrocytic uptake rate *τ*_au_ and transport *τ*_*g*_. Each data point represents the mean value obtained at the specified parameter values. Effects of varying *τ*_*g*_, i.e., recovery time of glutamate from astrocytes is shown in color-coded traces. Overlapped vertical and horizontal broken gray lines represent the approximate mean value of glutamate decay time (*τ*_decay_ analogue for astrocytic glutamate uptake, *τ*_au_) obtained from experiment (Figure 3C) and the approximate SB statistical mean values obtained from MEA data analysis respectively. Consistent to results with DHK experiments, as astrocytic uptake of glutamate is inhibited, viz. *τ*_au_) increases, (B-a) the firing rate also increases, (B-b) SB index decreases, (B-c) SB duration increases and (B-d) SB rate also increases. Moreover, the SB statistics were found to match with experimental observations closely at *τ*_*g*_ around 30s.

### SB can be produced with fixed amount of glutamate in astrocytes

*A* remains nearly constant during SBs for different values of *τ*_au_ as shown in Figures 7A, a–b. These results indicate that astrocytes do not directly take part in the generation of an SB. Their primary role is to restrict the amount of glutamate available for synaptic dynamics. By fixing *A* as a constant (Figures 7A, c–d), SBs can still be generated with features similar to the experiments, indicating that the dynamics of *A* may not be important for the generation of SB. By fixing *A*, the tripartite synapse in fact gets reduced into a bipartite synapse. As a bipartite (TUM) synapse is governed by *X* + *Y* + *Z* = 1, fixing *A* was equivalent to *X* + *Y* + *Z* = 1 − *A*, were 1 − *A* became another constant which is *<*1. This strongly suggests that the main role of astrocytes in generation of SB is to limit the amount of glutamate available in the bipartite synapse. The availability of glutamate then governs the firing patterns in SBs.

### Slower glutamate recovery from astrocytes in neurons alters SB properties

According to our TUMA model, the amount of glutamate in the *A* state can be varied either by varying *τ*_au_ or *τ*_*g*_. Here *τ*_*g*_ represents the timescale for glutamate-glutamine cycle which is followed by astrocytic glutamate uptake. Previous lines of research have shown the importance of this cycle for the generation of bursting activities in neurons (Bacci et al., 2002; Tani et al., 2014). Figure 7B shows network bursting statistics under combinations of different *τ*_au_ and *τ*_*g*_. Although *τ*_*g*_ has been used as an overly simplified representation of multiple processes that occur in the course of glutamine transport from astrocytes to neurons in this study, these simulation results provide an estimated range of *τ*_*g*_ within which this recovery may take place. From the data pooled from our cortical culture experiments, the reference SB statistics were found to be in the following ranges: array-wide firing rate (≈26 spikes/s), SB duration (≈0.29 s), SB index (≈0.82), and SB rate (≈5 SBs/minute). Also, the glutamate decay time (*τ*_decay_) measured across networks was found to be ≈0.26 s. In order to align with these values, it can be seen in Figure 7B that *τ*_*g*_ should lie around 30 s range. Also, the effect of increasing *τ*_*g*_ on bursting activities were in line with the effect of blocking astrocytic glutamate-glutamine cycle, as shown in the previous reports (Bacci et al., 2002; Tani et al., 2014).

## Discussions

Synchronous bursting (SB) is a network phenomenon and believed to be related to the proper or malfunction of the network. As the results shown above, the properties of the SB can be altered by local changes of glutamate dynamics in the level of individual synapses. Of particular importance is the GLT-1 GluTs expressed on the astrocytes which transport the majority of glutamate in the synapses into the astrocytes. In our model, the local changes in the transportation of glutamate in the synapses are manifested as the changes in the properties of the SB in the network. Therefore, the global changes of SB in the network level observed in our experiments can be used to infer useful information on the local glutamate transport; especially with the participation of the astrocytes in a tripartite synapse.

Although SB has been observed for a long time, its fundamental mechanism is still not clear. With our experiments and modeling, it is clear that the SB is the result of the interaction between the neuronal and glial systems in the network. In our TUMA model, the SBs originate from the asynchronous releases of glutamate from neurons and the effects of these random events get amplified by the positive feedback of the recurrent network in the culture as elevated firing spreading across the network. Our important finding is that the astrocyte and its glutamate handling properties are playing a crucial role of arresting this rapid network firing by providing a negative feedback in reducing the availability of glutamate in the neuronal system. The observed SB is then the result of the sequential manifestation of the positive and negative feedback in the neuronal and glial systems respectively in the network. Details of the SB mechanism in our TUMA model are described below.

### Initiation and positive feedback

In the dormant phase of the network, the number of asynchronous release increases (see Figure 2) before SB. These releases come from the noise-induced release of *X* which build up slowly during the dormant state. If these release events become large enough and the connectivity in the network is also high enough, action potentials gets triggered in some neurons. Once this happens, more glutamate gets released by these action potentials triggering more neurons to fire due to recurrent connections in the system. Consequently, this positive feedback then triggers a system-wide firing event.

### Maintenance of burst (active phase)

The positive feedback can be maintained as long as there is enough *X* to be released to trigger further action potentials. Depending on the system parameters (*A* level), sub-burst in the form of reverberations can also be possible because of the interaction between synaptic facilitation and depression in the TUM mechanism. One would expect reverberations to be more easily observed in a system with only excitatory inputs similar to experiments with bicuculline. This is because for systems with both inhibitory and excitatory inputs, there should be a wider and weaker distribution of input currents to the neurons. Intuitively, system with higher *X* (lower *A*) should have longer bursting duration because it will take longer for *X* to deplete to a certain threshold. Indeed, we see from both experiments (Table 1) and simulations (Figure 7A) that SB duration decreases with the amount of glutamate in the astrocyte.

### Negative Feedback and Termination of burst

During the active state of SB, the ready-to-release glutamate molecules (*X*) are constantly being transformed into *Z*, via multiple timescale processes, which will then be turned slowly back to *X* controlled by another long time constant. Due to overall slower recovery mechanism, *X* is not replenished fast enough from *Z*. The action potential triggered release from the pre-synaptic cell will be then too small to elicit action potentials at the post-synaptic cell. In this scenario, the positive feedback can no longer be sustained and the bursting will stop.

### Dormant phase (inactive phase)

Once the system is in the dormant state, system-wide firings stop and there are only isolated spikes created by noise-driven releases. During this phase, *X* can slowly recover from *Z*. As *X* increases, the amount of noise-driven-released glutamate will also increase, raising the probability of triggering a system-wide depolarization.

With the picture described above, the SB phenomenon can be considered as a generic emerging property arising from the negative and positive feedback interaction between the glial and neuronal networks. No special circuits (or buster neurons) or dis-inhibition (Figure 4E), the two traditional bursting mechanisms, are needed for its generation. The most important finding in our work is that, the firing and bursting patterns of the neurons are governed by the amount of glutamate in the astrocytes (*A*). It is possible that different regions of brain can have different amount *A* in the astrocytes locally, allowing different firing and bursting patterns at different brain regions. In this sense, the astrocytes are regulating the bursting property of neurons in the network.

As shown in the simulation results, the astrocytes maintain a much higher level of glutamate than neurons. This trafficking arises due to the simultaneously occurring fast synaptic glutamate uptake via its transporters and slow glutamate-glutamine cycle processes. This mismatch between the two timescales in effect results in accumulation of higher amount of glutamate in the astrocytes. Hence, the level of glutamate availability in the presynaptic neuron gets restricted. Presumably, for normal function of the neural network, the amount stored in or released from the astrocytes are adaptive to the problem at hand. A low astrocytic glutamate concentration is necessary for reliable synaptic transmission of information (Flanagan et al., 2018) and our case of high glutamate concentration in astrocytes during SB is consistent with the enhanced glutamate levels found in epilepsy. Arguably, the SBs observed in our experiments are similar to pathological states of the system.

Similar to the TUM model, it was assumed in our TUMA model that the amount of glutamate remains conserved within the tripartite synapse. During the different phases of SB, a very small change occurs in *A* (Figure 6C). In fact, if the amount of glutamate in the *A* state is fixed, qualitatively similar collective bursting patterns can still be obtained (Figure 7A). If *A* is fixed, we are back to the traditional bipartite synapse (a TUM model with *X*+*Y* +*Z* being a constant but less than one). Functionally, the astrocytes set the overall level of *A* through the two processes of uptake from *Y* and transforming *A* to *X*. Therefore, even if there is a violation of the conservation, it will not invalidate our conclusion as long as the overall *A* is fixed to a certain level.

During the active state, it is clear that the astrocytes exhibit SCEs (Figure 5). Its effects was not considered in our TUMA model. From Figure 5B, the SCEs in astrocytes were induced towards the termination of SBs. Longer SBs more reliably triggered SCEs in astrocytes than shorter SBs. This shows that astrocytic global calcium response depends on the amount of glutamate released in synapses. Interestingly, during the SCE transient in astrocytes, all the neurons maintained complete quiescence. Their firing activities recovered after the calcium decay. For such an observation, there could be multiple possible scenarios: First, calcium elevation may trigger astrocytic inhibitory gliotransmitters release into the synapses or extra-synaptic spaces which directly suppress the persisting neuronal activities (Montana et al., 2006). Two possible candidates are ATP/adenosine (Zhang et al., 2003; Panatier et al., 2011) and nitric oxide (Mehta et al., 2008) which have been reported to suppress hetero-synaptic activities. Second, the SCEs may also be associated with recruitment of additional astrocytic GluTs for more rapid uptake of glutamate from the synapse. These two processes may also occur simultaneously. Up till now, we have been discussing the effects of negative feedback of astrocyte. It is also known that astrocytic Ca elevation can enhance the excitability (facilitation) of the post-synaptic cells in the network (Verdugo et al., 2019). Concomitant astrocytic SCE response could be associated with positive feedback mechanism associated with increased glutamine transport to the neurons or calcium-induced glutamate release (Parpura and Haydon, 2000). It requires further more experiments to carefully dissect and confirm these speculations.

Finally, we like to point out that malfunction of astrocytic GLT-1 is related to neurological disorders such as seizure-like epilepsy (Tanaka et al., 1997; Werner et al., 2001) as well as behavioral disorders (Bechtholt-Gompf et al., 2010; Scofield and Kalivas, 2014; John et al., 2012). Moreover, electrophysiological evidence show that astrocyte-dependent glutamate recycling is important for active neurotransmissions (Tani et al., 2014). Our study in consolidation with previous reports clarifies the understanding of how altered uptake mechanisms or astrocyte-dependent glutamate recycling can affect neuronal network’s firings and collective behavior, which are representative of brain circuit’s functions. In the astrocyte targeted experiment of Bechtholt-Gompf et al. (2010) and John et al. (2012), a depression-like behavior was induced in rats by DHK treatment; the same drug used here. If we extend our findings to understand their observations, it seems that the increase in firings induced by the DHK are the origin of such changes. In terms of our model, this is just a decrease in the level of glutamate in the astrocytes (the *A* state). This last observation is consistent with the current view that astrocytes can be important in shaping the behavior of animals (Harada et al., 2016).

## Acknowledgements

This research work was supported by Taiwan’s MOST funds; 105-2112-M-001 -017 -MY3 and 108-2112-M-001 -029 -MY3.

